# Intermitochondrial signaling regulates the uniform distribution of stationary mitochondria in axons

**DOI:** 10.1101/2020.07.31.230250

**Authors:** Nozomu Matsumoto, Ikuma Hori, Tomoya Murase, Takahiro Tsuji, Seiji Miyake, Masaru Inatani, Yoshiyuki Konishi

**Author notes:** Corresponding author: Yoshiyuki Konishi, Department of Applied Chemistry and Biotechnology, Faculty of Engineering, University of Fukui, 3-9-1 Bunkyo, Fukui 910-8507, Japan.

## Abstract

In the central nervous system, many neurons develop axonal arbors that are crucial for information processing. Previous studies have demonstrated that premature axons contain motile and stationary mitochondria, and their balance is important for axonal arborization. However, the mechanisms by which neurons determine the positions of stationary mitochondria as well as their turnover remain to be elucidated. In this study, we investigated the regulation of spatiotemporal group dynamics of stationary mitochondria. We observed that the distribution of stationary mitochondrial spots along the unmyelinated and nonsynaptic axons is not random but rather relatively uniform both in vitro and in vivo. Intriguingly, whereas the positions of each mitochondrial spot changed over time, the overall distribution remained uniform. In addition, local inactivation of mitochondria inhibited the translocation of mitochondrial spots in adjacent axonal regions, suggesting that functional mitochondria enhance the motility of neighboring mitochondria. Furthermore, we showed that the ATP concentration was relatively high around mitochondria, and treating axons with phosphocreatine, which supplies ATP, reduced the immobile mitochondria induced by local mitochondrial inhibition. These observations indicate that intermitochondrial interactions, mediated by ATP signaling, control the uniform distribution of axonal mitochondria. The present study reveals a novel cellular system that collectively regulates stationary mitochondria in axons.

## INTRODUCTION

The regulation of mitochondrial transport in axons plays critical roles in the function of neurons by regulating axonal morphology and modulating presynaptic functions (1–6). They are frequently found in places consuming high energy, such as the nodes of Ranvier, presynaptic sites and growing axonal terminals (7–9). Microtubules and kinesin-1 play roles in transporting mitochondria from the cell body to axonal terminals, whereas dynein mediates retrograde transport. Defects in these transport systems are implicated in neurological disorders (10, 11). Despite the accumulating body of knowledge on the molecular mechanisms that regulate mitochondrial transport, the intracellular systems by which neurons simultaneously regulate mitochondrial distribution and dynamics along axons remain to be elucidated. Previous studies have revealed that in the axons of mature cortical or hippocampal neurons, mitochondria are immobilized for an extended period of time at presynaptic sites (9, 12).

Importantly, even in premature axons before synaptic maturation (i.e., 3-7 days in vitro (DIV)), mitochondria are generally stationary. The mitochondrial anchoring protein syntaphilin (SNPH) mediates mitochondria docking on microtubules (1, 13). Loss of SNPH dramatically increases the number of motile mitochondria in axons at early stages (1). Inhibition of SNPH function was accompanied by a reduction in axonal branches in cortical neurons at 5 DIV (2), indicating that these stationary mitochondria are required for the development of normal axonal arbors. Considering that axonal remodeling and presynaptic elimination take place in the adult central nervous system (CNS), these mechanisms may also contribute to the function of the CNS throughout one’s lifespan. Moreover, parts of the axonal segment do not contain a presynaptic structure or myelinated region, including the retinal nerve fiber layer, a typical example for unmyelinated and nonsynaptic axonal regions, in which mitochondrial movement is occasionally observed (14) to be different from the presynaptic site of cortical neurons (9). In addition, a recent study revealed that SNPH inhibits the degeneration of demyelinated axons (15). These evidences points to the importance of understanding the systems regulating the dynamics and distribution of stationary mitochondria in the absence of mature presynaptic sites or nodes of Ranvier. More than a decade ago, Miller and Sheetz reported (16) a uniform mitochondrial distribution along the axon of sensory neurons, although the generality of this important finding has not been verified, and its control mechanism is unknown. The present study aimed to uncover the system that regulates the distribution and dynamics of mitochondrial populations in axons.

## RESULTS

### Dynamics of axonal mitochondria in premature CGNs in vitro

We began our investigation to determine whether there is a system that regulates the distribution and/or group dynamics of axonal mitochondria by using young cerebellar granule neurons (CGNs). They do not form functional synapses with each other; thus, they are suitable for studying cell-autonomous systems. Although a large part of presynaptic sites remain orphaned, presynaptic clusters on axons start to make contact with postsynaptic sites around the first week in culture (17). To exclude the effect of postsynaptic sites, neurons were placed inside of a silicon chamber; the next day, the silicon chamber was removed to allow axons to grow outside of the plating area without contacting the dendrites of other neurons (Fig. 1A). After axonal extension, mitochondria were stained with MitoTracker Red CMH_2_-XRos. In accordance with previous reports of hippocampal neurons and cortical neurons (1, 2), both stationary and motile mitochondria exist in axons of CGNs at 3 DIV (Fig. 1B, C). The proportion of stationary mitochondria significantly increased as the axon matured, and most of the axonal mitochondria (93 ± 2%, n = 5 axons) were stationary for 15 min at 7 DIV (Fig. 1C, D).

**Fig. 1.**
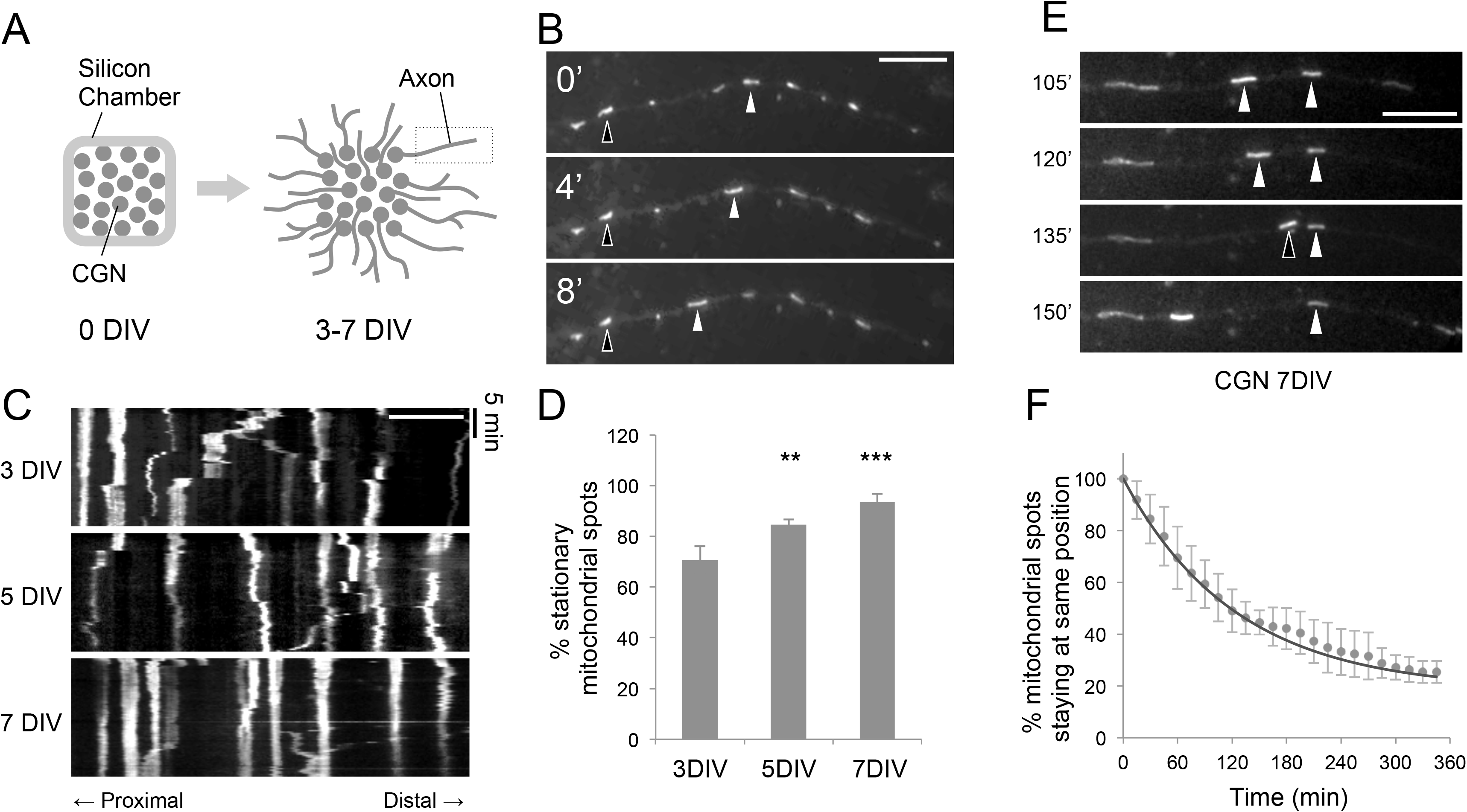
Dynamics of axonal mitochondria of premature CGNs in vitro. (A) Schematic presentation of the culture method to separate axons from somatodendrites. CGNs were plated inside of a silicon chamber that was attached to the dish. The silicon chamber was removed at 1 DIV to allow axonal extension. (B) Representative time-lapse images of an axon at 3 DIV stained with MitoTracker. Examples for motile (white arrowheads) and stationary (open arrowheads) are indicated. (C, D) Kymographs (C) and quantified data (D) showing progressive reduction of mitochondrial motility with axonal maturation (n = 5 axons in each condition). (E) Representative time-lapse images of the CGN axon at 7 DIV stained with MitoTracker. Images were taken at 15 min intervals, and those at the indicated time points are shown. White arrowheads indicate mitochondria that remained in the same position, whereas an open arrowhead indicates a mitochondrion that changed position. (F) The percentage of mitochondria in which the positional change was less than 5 μm from the initial position was quantified. The fitting curve using a one-phase decay model was also revealed. Data were obtained from n = 6 axons. Scale bars indicate 10 μm. Values represent the mean ± 95% CI, **p < 0.01, ***p<0.001, Tukey’s test.

We next asked whether those stationary mitochondria stay at the same position for an extended period of time. A previous study using mature cortical neurons revealed that stable mitochondria remain at synaptic sites over 12 hrs (9). We monitored axonal mitochondria of 7 DIV CGNs for 6 hrs in 15 min intervals. The mitochondrial spots that did not change location during the first 15 min were defined as stationary. In many cases, stationary mitochondrial spots paused at the same position for 15 min but occasionally changed their location (Fig. 1E). We quantified the percentage of mitochondrial spots that stayed at the same position from the beginning of the observational period and found that it decayed at a constant rate over time. Under this condition, 82% of stationary mitochondrial spots translocated at 96 min of half-life (t1/2), which was obtained by mathematically fitting the data to the one-phase decay model (Fig. 1F) (n = 6 axons). These results suggest that most stationary mitochondria become motile mitochondria in a relatively short time that is constant in premature axons. The model fitting indicated that a small population (18%) of stationary mitochondrial spots would not change their location for an extended period of time at 7 DIV. They likely represent immobilized mitochondria trapped in specific axonal sites, such as those in the mature presynaptic sites (9, 18).

### Stationary mitochondria distribute uniformly in axons

Next, we analyzed the spatial distribution pattern of stationary mitochondria. CGNs were cultured and labeled with MitoTracker as in Fig. 1, fixed at 7 DIV, and stained with tubulin antibody to visualize axonal processes (Fig. 2A). Images of axonal segments (0-120 μm from the terminal) that do not have branches were analyzed. After linearization and binarization, the number of mitochondrial spots in each compartment was quantified, and the Iδ-index, a dispersion index that enables the distribution pattern to be identified, was calculated. If the spatial distribution is random, the value of Iδ will approximately 1. If the spatial distribution is uniform, Iδ will be less than 1, and it will approach 1 as the compartment size increases. For a clustered distribution, Iδ will become more than 1 (Fig. 2B) (19). We found that the distribution of mitochondrial spots is typically uniform (Fig. 2C). Likewise, the frequency distribution of mitochondria was significantly different from the Poisson distribution, which represents a random distribution (Fig. 2D) (at 20 μm compartment size; p < 0.01 in χ^2^ analysis, n = 20 axons). The density of mitochondrial spots in CGN axons at 7 DIV was 10.3 ± 0.7 per 100 μm axonal segment. Given that most of the mitochondria of CGN axons are stationary at 7 DIV, our observations indicate a uniform distribution of stationary mitochondria.

**Fig. 2.**
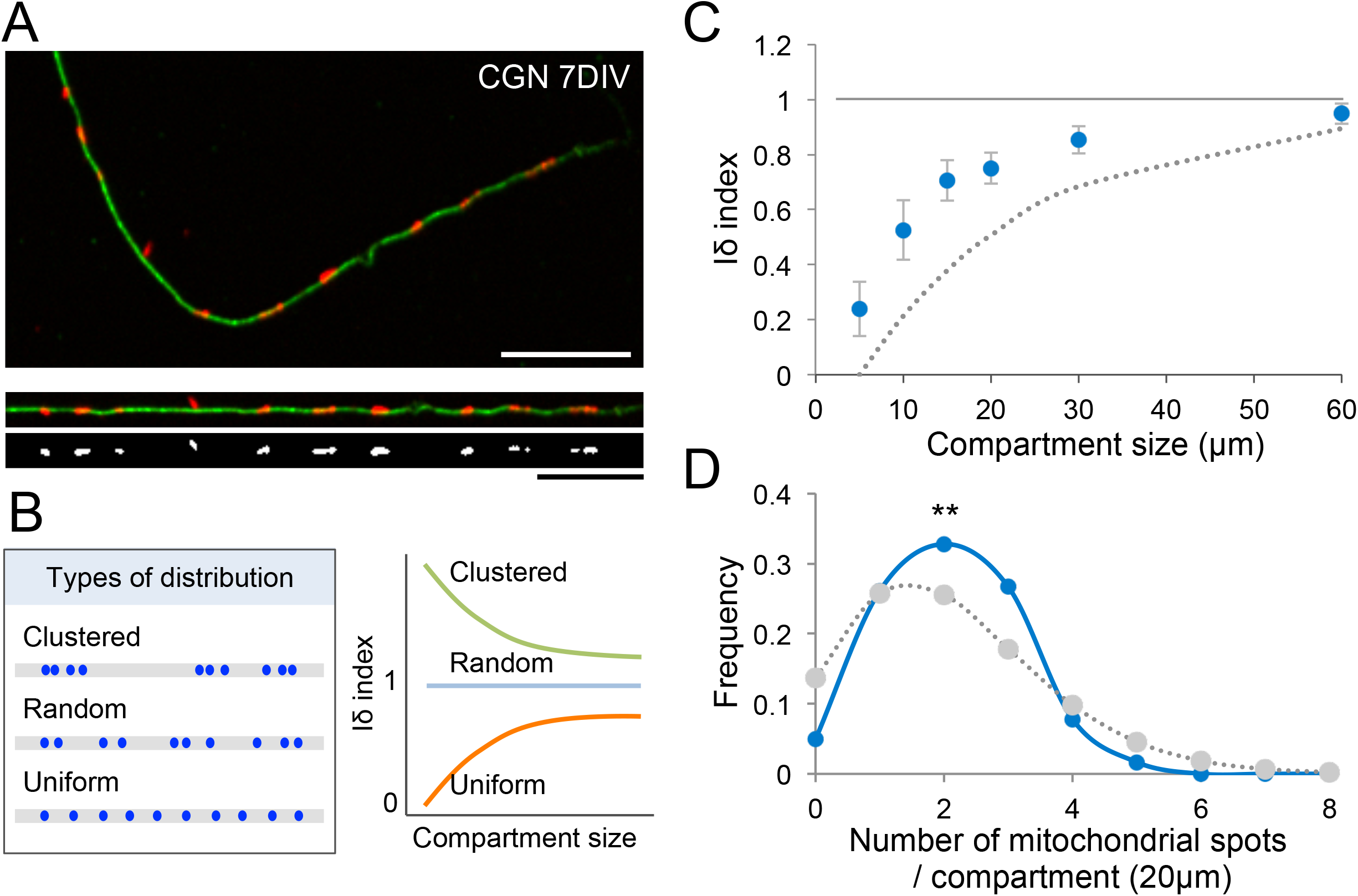
The distribution of mitochondrial spots along axons is relatively uniform in vitro. (A) Representative images of a 7 DIV CGN axon stained with MitoTracker and anti-tubulin antibody. Examples of image processing for compartment analysis showing straightened axons and binary images are also depicted at the bottom. Scale bars indicate 20 μm. (B) Schematic presentation of typical distributions and Iδ-index profiles of each case. (C) The Iδ-index profile of axonal mitochondrial spots obtained from cultured CGN axons at 7 DIV. Data were obtained from distal axonal segments (0 - 120 μm from the terminal, n = 20 axons). Values represent the mean ± 95% CI. The dotted line indicates the Iδ-index profile for complete uniform distribution. (D) Frequency of observed mitochondrial spots (solid line) versus Poisson distribution (dotted line) at a compartment size of 20 μm indicated that the mitochondrial spot distribution is significantly different from a random distribution (**p < 0.01, χ^2^ analysis).

We also analyzed the mitochondrial distribution in axons of cultured retinal ganglion cells (RGCs) at 8 DIV. Positions of stationary mitochondria were determined from the kymograph of time-lapse images, since a considerable number of motile mitochondria were observed in cultured RGC axons (Fig. S1A). Quantification revealed a relatively uniform distribution of stationary mitochondria in RGC axons (Fig. S1B, C).

### Mitochondrial distribution is relatively uniform in CNS axons in vivo

To address whether mitochondria are uniformly distributed in vivo, we introduced expression vectors for mCherry-Mito and enhanced green fluorescent protein (EGFP) to the cerebellum of postnatal day 6 (P6) mice by electroporation (20). To minimize the effect of postsynaptic regulation, the cerebellum was isolated at P10, in which parallel fiber-Purkinje cell synapses have not yet been established (21). In coronal sections, EGFP-positive parallel fibers were detected from the inner part of the external granular layer to the molecular layer (Fig. 3A). The distribution of mitochondrial spots along parallel fibers was analyzed as shown in Fig. 2. The density of mitochondrial spots in parallel fiber axons of P10 mice was 4.8 ± 0.5 per 100 μm, and some axons revealed high mitochondrial density. Clustering of mitochondrial spots was detected occasionally, as revealed by an increased Iδ-index at a compartment size of 10 μm (Fig. 3B). Nevertheless, the Iδ-index profile revealed that mitochondrial spots are relatively uniformly distributed (Fig. 3B), and the frequency distribution of mitochondria was significantly different from the Poisson distribution (at 40 μm compartment size; p < 0.05 in χ^2^ analysis, n = 22 axons), as it had a higher value around average density (Fig. 3C).

**Fig. 3.**
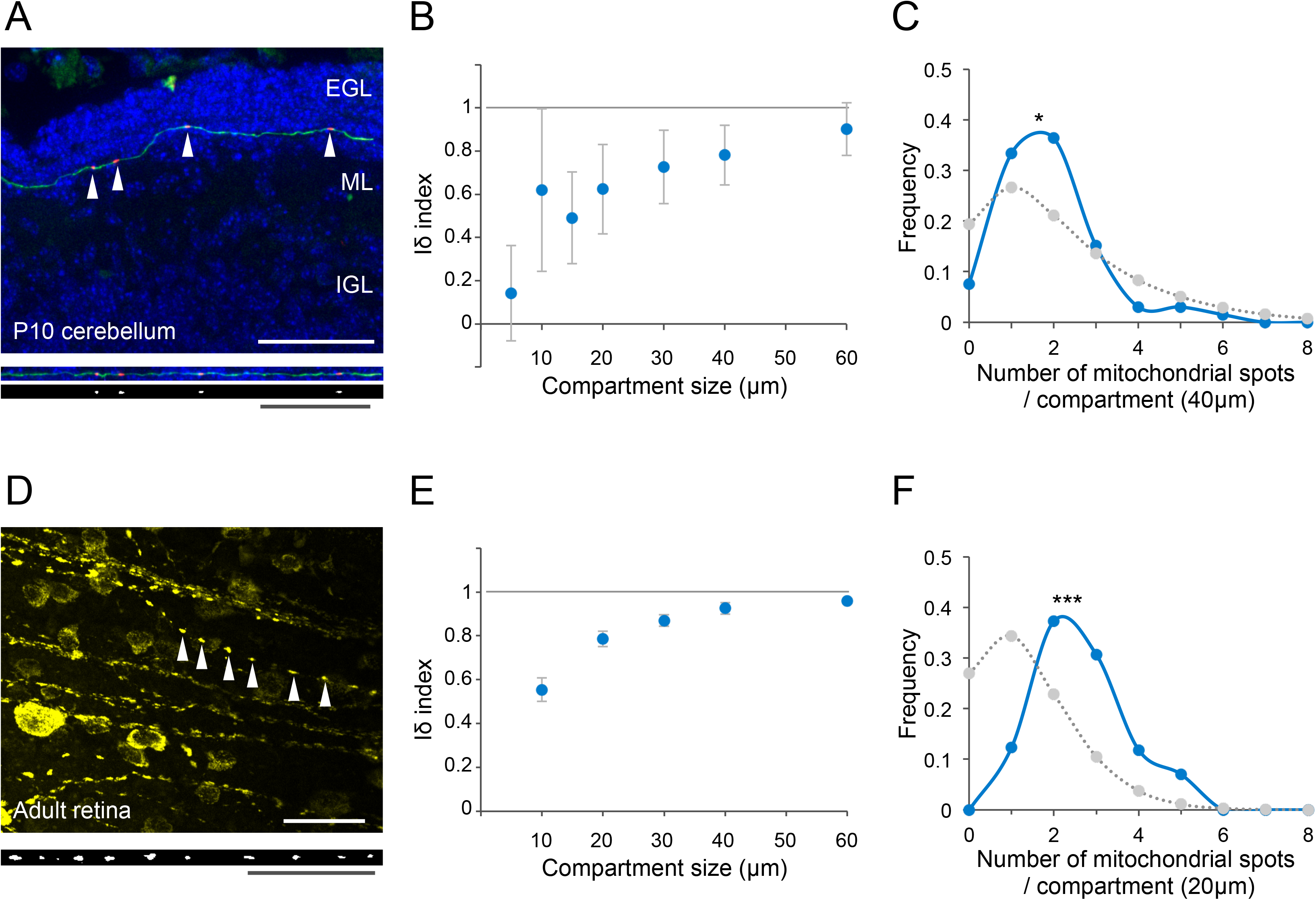
Distribution of mitochondrial spots in CNS axons in vivo. (A) Representative image of a coronal cerebellar section from a P10 mouse, which has been electroporated with expression vectors for mCherry-Mito (Red; arrows) and EGFP (Green) at P6. Layer structure of the cerebellum visualized by nuclear staining (blue) indicating that the CGN axon extends in the external granule layer. Straightened binary images indicating mitochondrial position are also shown (bottom). EGL: external granule layer, ML: molecular layer, IGL: internal granule layer. (B) Iδ-index profiles of axonal mitochondrial spots in CGN axons obtained from P10 cerebella. Data were collected from n = 22 axons (120 μm segment) from 4 cerebella. (C) Frequency of mitochondrial spots observed in vivo (solid line) versus a Poisson distribution (dotted line). Since the mitochondrial density was lower than in vitro culture, data were analyzed by 40 μm of compartment. (D) Representative image as well as straightened and binary images of RGC axons in the retinal nerve layer of *Thy1-mitoYFP* transgenic mice. The axonal region where mitochondria (arrows) within a single axon can be visualized was used for the analysis. (E) Iδ-index profiles of axonal mitochondrial spots in RGC axons obtained from 12-week-old mouse retinas. Data were collected from n = 38 axons (120 μm segment) from 5 animals. (F) Frequency of mitochondrial spots observed in retina (solid line) versus a Poisson distribution (dotted line). Data were analyzed by 20 μm compartments. Values represent the mean ± 95% CI (*p < 0.05, ***p < 0.001, χ^2^ analysis). Scale bars indicate 50 μm.

Since RGC axons in retinal nerve fibers are unmyelinated and do not form synapses even in the adult stage, we also analyzed the mitochondrial distribution in this region. We generated transgenic mice that express mitochondria-targeted yellow fluorescent protein (YFP) under the control of the *Thy1* promoter (*Thy1-mitoYFP*). Flat-mounted retinas prepared from 12-week-old mice revealed a large number of YFP-positive RGCs throughout the retina (Fig. S2) and YFP-positive mitochondria in RGC axons (Fig. 3D). The density of mitochondrial spots in RGC axons was 13.1 ± 0.4 per 100 μm. The Iδ-index profile revealed that mitochondrial spots were relatively uniformly distributed (Fig. 3E), and the frequency distribution of mitochondria was significantly different from the Poisson distribution (at 20 μm compartment size; p < 0.001 in χ^2^ analysis, n = 38 axons) (Fig. 3F). In conjunction with primary culture neuron analysis, these results suggest the existence of systems that uniformly distribute axonal mitochondria in vivo.

### Positions of stationary mitochondria change over time

These observations led us to hypothesize two possible mechanisms for regulating the distribution of stationary mitochondria (Fig. 4A). In the first model (a), mitochondrial anchoring sites are predetermined by anchoring molecules or other factors uniformly distributed in axons. In this case, the overall distribution of stationary mitochondria does not change over time, since even if a stationary mitochondrion moves away from a particular site, another mitochondria will be anchored at the same site. In the second model (b), anchoring sites for mitochondria are not predetermined, whereas mitochondria communicate with each other to control distances. In this case, the distribution of stationary mitochondria will change over an extended period of time. We noticed that the distribution of axonal mitochondria mostly changed in 120 min (Fig. 4B), by which time the majority of stationary mitochondria were replaced (Fig. 1F). We extracted the positions of mitochondria that stayed at the same position between two adjacent frames (interval = 15 min). When these mitochondrial positions were compared with the positions of stationary mitochondria at 120 min, Pearson’s correlation coefficient was largely decreased (r = 0.32, p = 0.44) and remained low after 240 min (r = 0.41, p = 0.32) (Fig. 4C). In contrast, the Iδ-index for the distribution of stationary mitochondria did not markedly change with time (e.g., 0 min; 0.76 ± 0.04, 120 min; 0.73 ± 0.04, 240 min; 0.70 ± 0.05, at 20 μm of compartment size) (Fig. 4D). These results suggest that mitochondrial positions are not predetermined, but their relative position is maintained.

**Fig. 4.**
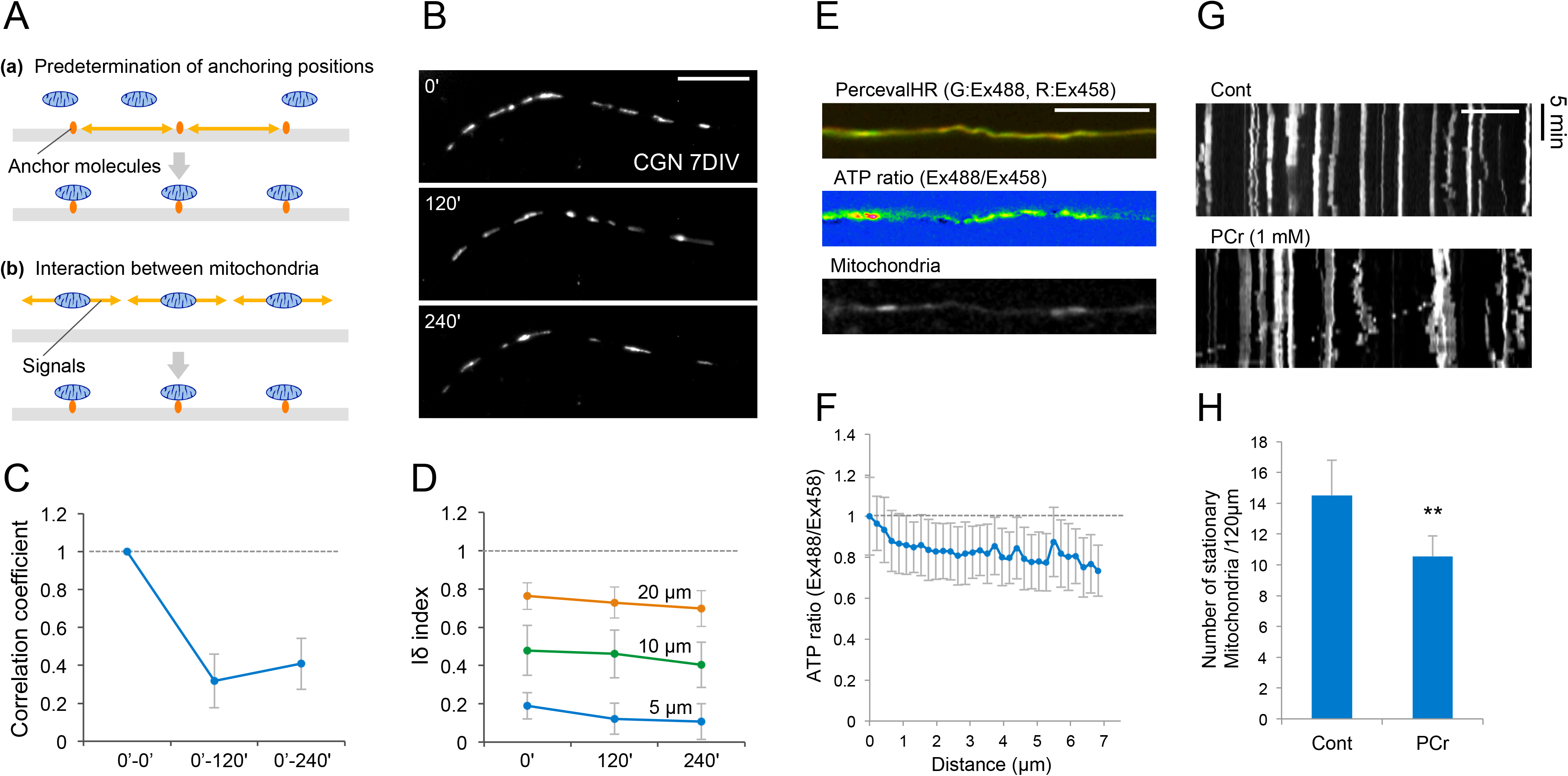
Analysis of the mechanisms that uniformly distribute mitochondria in axons. (A) Schematic presentation of two models to control uniform mitochondrial distribution; anchoring positions were predetermined (a) or determined by intermitochondrial interaction (b). (B) Representative images of axonal mitochondria at the indicated time points. Note the difference in mitochondrial positions at different times. Scale bar indicates 20 μm. (C) Pearson’s correlation coefficient of mitochondrial position between the indicated time points revealed a large reduction in 120 min. Data were obtained from n = 8 axons. (D) Iδ-index of axonal mitochondrial spots at three different compartment sizes showing that mitochondrial distribution is relatively uniform over time. (E) Axon of CGN expressing the ATP sensor PercevalHR and mCherry-Mito. Signals for ATP (Ex 488), the isosbestic point (Ex 458) and the ATP ratio (Ex 488/458) are shown. Scale bar indicates 10 μm. (F) The ATP ratio in each position relative to the terminal of the mitochondrial position was revealed. Axonal regions with distances between mitochondria ranging from 10 to 15 μm were analyzed (n = 44 axonal regions). (G, H) Representative kymographs (G) and quantified results (H) of axonal mitochondria in the presence or absence of 1 mM phosphocreatine (PCr) (control; n = 16 axons, PCr; n = 22 axons in 4 experiments). Images were taken for 15 min. Scale bar indicates 20 μm. Values represent the mean ± 95% CI. **p < 0.01, Welch’s t-test.

### Function of ATP on the regulation of stationary mitochondria

If mitochondria communicate with each other to maintain a uniform distribution, some signaling molecules should be present. Since mitochondria produce ATP, clustering of stationary mitochondria may increase the local ATP concentration. Therefore, we investigated whether ATP itself functions as a signaling molecule to regulate the distribution of stationary mitochondria. First, we expressed PercevalHR, a genetic sensor for detecting the ATP:ADP ratio (22) in CGN. It senses changes in low concentrations of ATP compared to the original Perceval concentration. Signals excited by a 488 nm (equivalent to ATP) laser were normalized by signals excited by 458 nm (close to the isosbestic point). Treatment with 1 μM of oligomycin, an inhibitor of mitochondrial ATP synthase, significantly reduced PercevalHR (ex488 nm/ex458 nm) signals (48 ± 1% relative to the control) in CGN (Fig. S3). Reduction of PercevalHR signals was also detected to a lesser extent by administration of 50 mM 2-deoxyglucose (86 ± 1%). Even though there was a large axon-to-axon variation, PercevalHR signals were gradually decreased depending on the distance from the mitochondria (Fig. 4E, F). Notably, there was a limitation that affected the measurement of absolute ATP concentrations. Nevertheless, these results suggest that relative ATP concentrations tend to be low at the axonal segment away from mitochondria, indicating that the density of stationary mitochondria could alter local ATP concentration.

Second, we asked whether ATP has a role in regulating the distribution of stationary mitochondria. To this end, 7 DIV CGNs were treated with phosphocreatine, which provides high-energy phosphate to convert ADP to ATP and is used to supply ATP to cells, including neurons (23, 24). Time-lapse imaging of axonal mitochondria revealed that phosphocreatine treatment increased the motility of mitochondria and significantly reduced the number of stationary mitochondria (Fig. 4G, H, 14.5 ± 1.2 in controls versus 10.5 ± 0.7 in phosphocreatine treatment per 100 μm of axonal segment, p <0.01, Welch’s t-test).

### Local inactivation of mitochondria affects the translocation of neighboring mitochondria

The results described above support the notion that the uniform distribution of stationary mitochondria is regulated by intermitochondrial signaling. To substantiate this idea, it would be desirable to control the signal molecule at specific axons, but it is technically difficult to locally manipulate the concentration of signal molecules such as ATP. Hence, we decided to locally manipulate the function of mitochondria instead by chromophore-assisted light inactivation (CALI) (Fig. 5A). It has been shown that light irradiation of mitochondria-localized KillerRed (KillerRed-dMito) results in the production of reactive oxygen species (ROS), thereby inactivating mitochondria locally at an illumination site (3, 25). Local illumination of green light resulted in the bleaching of KillerRed-dMito at a specific axonal site (Fig. 5B). Subsequently, the positions of unilluminated mitochondria around the illumination site were monitored (Fig. 5C). To minimize KillerRed-dependent phototoxicity, images of both proximal and distal sites (up to 50 μm from the border of the illumination site) were taken at 0, 30 and 60 min after illumination (Fig. 5C). The number of immovable mitochondria was quantified as shown in Fig. 1F. In control axons that did not receive photoillumination or that expressed mCherry-Mito instead of KillerRed-dMito, most of the mitochondria changed their location in 60 min (immovable mitochondria; 45.0 ± 5.0% in KillerRed-dMito-expressing neurons without photoillumination, 38.7 ± 4.8% in mCherry-Mito-expressing neurons with photoillumination) (Fig. 5C, D). In CALI-applied axons, the rate of immovable mitochondria was significantly increased (63.4 ± 2.5%) compared with that in controls (Fig. 5D; p < 0.05 compared with no illumination, p < 0.01 compared with mCherry-Mito, Tukey’s test). Furthermore, phosphocreatine treatment prior to CALI application reduced the rate of immovable mitochondria (38.4 ± 4.5%, p < 0.001 compared with CALI-treated axons, Tukey’s test). These results indicate that mitochondrial function increases the motility of other mitochondria and that ATP contributes to intermitochondrial signaling.

**Fig. 5.**
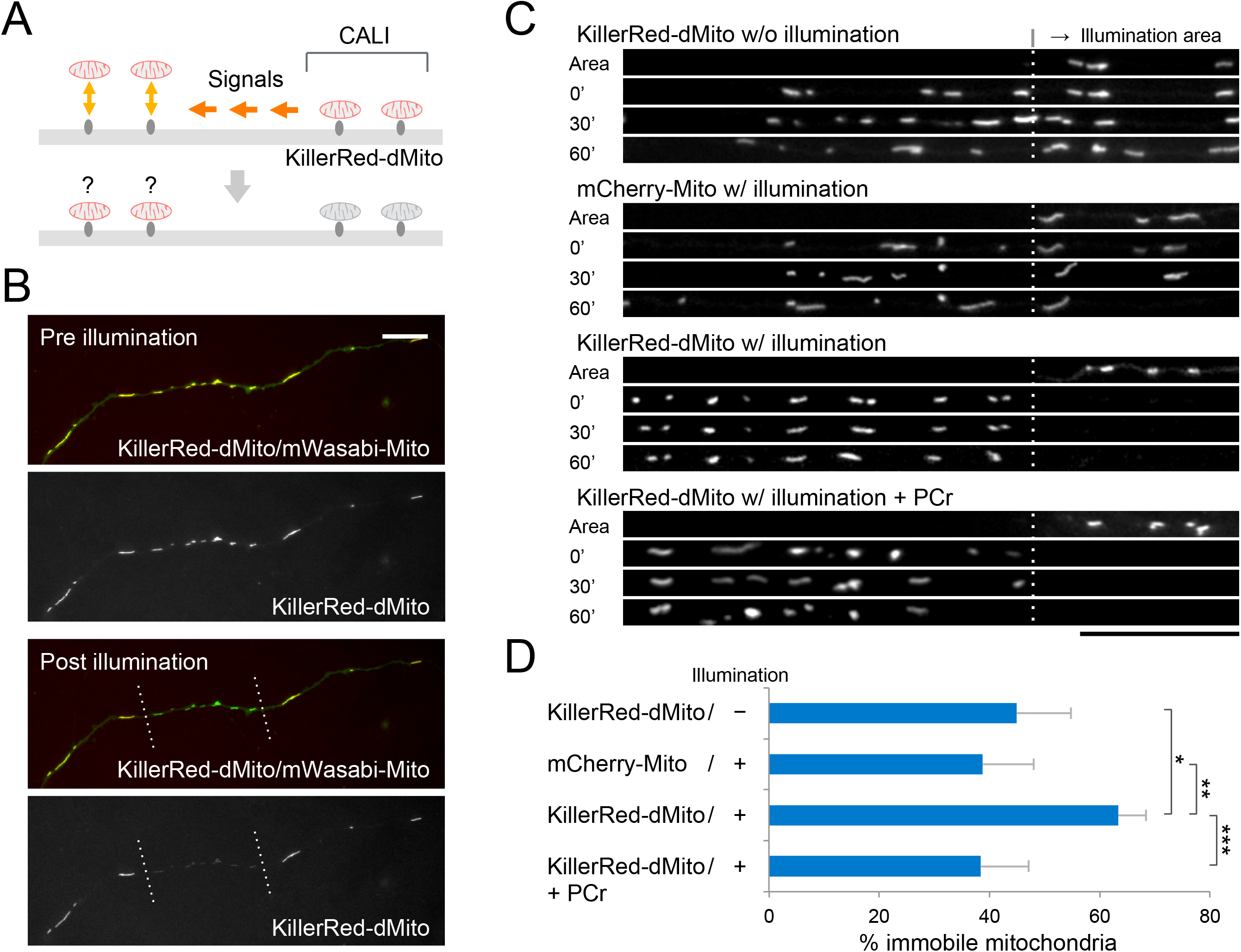
Effect of local mitochondrial inactivation on mitochondrial dynamics. (A) Schematic presentation of the procedure for CALI-mediated analysis of intermitochondrial interactions. Mitochondria were locally inactivated by CALI, and the dynamics of neighboring mitochondria were analyzed. (B) Representative images of axonal mitochondria expressing Mito-KillerRed before and after local illumination. Signals for mWasabi-Mito were also revealed. (C) Representative images of linearized axons at the indicated times after photoillumination. Part of the illumination areas (Area) and proximal regions to the illumination area (up to 50 μm from the border) are shown. First row: neurons were expressed with Mito-KillerRed, but no illumination was applied Second row: neurons expressing mCherry-Mito as a control were locally illuminated. Third row: neurons expressing Mito-KillerRed were locally illuminated. Fourth row: neurons expressing Mito-KillerRed were pretreated with phosphocreatine (PCr) before the illumination. (D) Percentages of immobile mitochondria 60 min after CALI were measured as in Fig. 1F. Local inactivation of mitochondria significantly increased the ratio of immobile mitochondria at the neighboring region, and phosphocreatine was reduced to the control level. In each condition, n = 12 axons in 3 experiments were analyzed. Scale bars indicate 20 μm. Values represent the mean ± 95% CI, *p < 0.05, **p < 0.01, ***p<0.001, Tukey’s test.

## DISCUSSION

In this study, we showed that the distribution of stationary mitochondrial spots in premature CGN axons is not random but rather relatively uniform both in vitro and in vivo. Similar results were obtained from RGC culture and adult RGCs in the retina. These observations reaffirm the previous finding in cultured sensory neurons (16) and suggest that neurons generally possess a system that uniformly distributes mitochondria along their axons. In contrast to the presynaptic site of mature cortical neurons, in which mitochondria are immobilized over several hours (9), stationary mitochondria changed their location in a relatively short time in premature CGNs (t_1/2_ = 96 min, at 7 DIV). Intriguingly, the positions of mitochondria changed over time, suggesting that the sites for mitochondrial capture changed over time.

What is the meaning of the uniform mitochondrial distribution? A previous study demonstrated that a reduction in stationary mitochondria in premature cortical neurons causes a reduction in axonal branches (2, 26). Thus, stationary mitochondria are required for establishing and/or maintaining proper axonal arbors, likely by locally supplying ATP and/or regulating Ca^2+^ concentrations. If distribution is random, it is likely that mitochondrial deficiency will occur stochastically in certain axonal regions. In addition, premature axonal arbors undergo dynamic morphological changes, and stationary mitochondria need to be provided in newly growing branches. Constantly switching between motile and stationary states could enable mitochondria to adapt to such situations. Since axonal arbor morphology becomes static as an axon matures, neurons might regulate the balance between motile and stationary mitochondria depending on axonal maturation.

The present study revealed that axons possess a system that regulates mitochondrial distribution via intermitochondrial signals for the following reasons. If mitochondrial distribution is determined by the preexisting structure, such as presynaptic sites, the distribution of mitochondria should not largely change over time because after a mitochondrion moves away, another mitochondrion should be captured at the same position. However, our observations revealed that the positions of each mitochondrion changed considerably, suggesting the possibility that mitochondria remain distant from each other. Consistently, locally inhibiting mitochondria by applying CALI prevented the translocation of mitochondria at the neighboring region, revealing that functional mitochondria enhance the motility of other mitochondria. These observations are consistent with a previous report that mitochondria tend to be captured between preexisting mitochondria (16).

Producing ATP and controlling the Ca^2+^ concentration are the main functions of mitochondria. Our results suggested that ATP mediates intramitochondrial signaling, although we do not exclude the possibility of the contribution of other signaling molecules, including Ca^2+^, that have been reported to affect mitochondrial motility (27–29). We demonstrated that supplying ATP with phosphocreatine application reduced immobile mitochondria that were induced by local mitochondrial inactivation. Since ATP is a small molecule, it should be rapidly diffused in the axoplasm after being produced by mitochondria, but rapid consumption in the axonal cytoplasm may create an ATP gradient. Indeed, in a previous study, an ATP gradient along the axon was observed depending on the distance from the growth cone (30). Using an ATP censor, we detected ATP concentration differences depending on the distance from mitochondria. This result contradicts a previous study reporting that ATP is uniformly distributed along axons (31). We believe that this discrepancy could be the result of a couple of differences. First, the PercevalHR that we used in the current study is more sensitive than the previously used Perceval (22). Second, while a previous study also reported ATP fluctuations in axons, the position of mitochondria was not taken into account since analysis focused on the ATP concentration throughout the axon. As mitochondria are distributed uniformly along axons, they will not be detectable unless they are compared with the distance from mitochondria in detail.

Our present study revealed an intracellular system regulating mitochondrial distribution and dynamics in axons. This system likely plays important roles in axonal arborization and reorganization. Finally, it is also interesting to note that axotomy induces a local reduction in mitochondrial motility accompanied by ATP depletion, which restricts axonal regeneration (30). Studying the molecules that receive ATP signals in the future could be valuable as it could lead to the ability to achieve efficient axonal regeneration.

## MATERIALS AND METHODS

### Primary culture of neurons

All animals were treated according to the institutional ethical guidelines, and experiments were approved by the animal ethics committees of the University of Fukui. Primary cultures of mouse CGNs were prepared from ICR (Jcl:ICR) mice (CLEA Japan, Inc, Tokyo, Japan) at postnatal days 4 to 6 as described previously (32). CGNs were dissociated with trypsin and plated on glass (at 0.25-0.5 × 10^6^ cells/cm^2^, depending on the experiment) that had been coated with poly-L-ornithine and attached to a silicon chamber (flexiPERM; Sarstedt, Nümbrecht, Germany). CGNs were maintained in Basal Medium Eagle (BME; Sigma-Aldrich, St. Louis, MO) supplemented with 10% calf serum (Thermo Fisher Scientific Inc., Waltham, MA), penicillin (1 mg/ml), streptomycin (1 mg/ml), glutamine (2 mM) and KCl (25 mM). For time-lapse analysis, Minimal Essential Medium without phenol red (MEM; Thermo Fisher Scientific Inc.) supplemented with 20 mM Hepes-KOH pH 7.2 was used instead of BME. At 1 DIV, cytarabine was added to the culture (10 μM), and the silicon chamber was removed to visualize the axonal extension.

### Transfection in primary culture neurons

To introduce plasmids into cultured CGNs, either electroporation or the calcium phosphate method was used. Electroporation was performed as previously described (33). CGNs (3 × 10^6^ cells) and plasmids were suspended in 70 μl of Dulbecco’s modified Eagle’s medium (Sigma-Aldrich) and exposed to decay pulses using a CUY21-edit II pulse generator (BEX, Tokyo, Japan) as follows: poring pulse, 275 V for 1 ms; driving pulse,+ 20 V for 50 ms × 5 times at 50 ms intervals with reversal of polarity. In the experiment shown in Fig. 5, 1.2 μg of KillerRed-dMito (Evrogen, Moscow, Russia) or mCherry-Mito-7 (Addgene #55102) (34) together with 2.8 μg of mWasabi-Mito-7 (Addgene #56508) (35) and 0.5 μg of bcl-xl/pcDNA3 (20) was introduced. Immediately after electroporation, prewarmed media was added to the cell, and 5 × 10^5^ cells were spread in the grass bottom plate attached with flexiPERM midi (Sarstedt). The calcium phosphate method was performed as previously described (32). Prior to transfection, neurons were incubated in Dulbecco′s Modified Eagle′s Medium (DMEM; FUJIFILM Wako Pure Chemical Co., Osaka, Japan) at 37 °C in a CO_2_ chamber. For the experiments shown in Fig. 4E, 3 μg of pGW-PervevalHR (Addgene #57432) (22), 0.1 μg of mCherry-Mito-7 and 0.8 μg of bcl-xl/pcDNA3 were suspended in 40 μl of 250 mM CaCl_2_ solution mixed with the same amount of 2×HBS solution (270 mM NaCl, 9.5 mM KCl, 1.4 mM NaH_2_PO_4_, 15 mM glucose, 42 mM Hepes, pH 7.1). After 15 min, the mixture was added to the culture and incubated for 15 min in a CO_2_ incubator. After washing twice with DMEM, neurons were placed in the conditionalized medium.

### Cell imaging

To stain axonal mitochondria, neurons were treated with 500 nM MitoTracker Red CM-H_2_XRos (Thermo Fisher Scientific Inc.). For immunocytochemistry, neurons were fixed for 20 min with 4% paraformaldehyde (PFA) in phosphate buffered saline (PBS). Following permeabilization with 0.4% Triton X-100/PBS for 15 min, neurons were placed in blocking solution (5% goat serum, 3% bovine serum albumin and 0.02% Tween 20 in PBS) and then treated with a monoclonal antibody against α-tubulin (12G10 at 1:1000; Developmental Studies Hybridoma Bank of University of Iowa) in blocking solution at 4 °C overnight. Goat anti-mouse IgG conjugated to Alexa Fluor 488 (1:1000; Abcam, Cambridge, UK) was the secondary antibody. For live-cell imaging of CGNs, a glass bottom dish was placed in a stage top incubator (ZILCS; Tokaihit, Shizuoka, Japan) maintained at 37 °C with a supply of 5% CO_2_. Mitochondria in the CGN axon were observed by using an Axiovert 200 M equipped with an MRm monochromatic digital camera (Carl Zeiss, Oberkochen, Germany). Images were acquired with 1388 × 1040 pixels using a 40× objective lens. For the CALI technique, the results for which are shown in Fig. 5, a small circular area defined by an iris was illuminated for 30 s with orange light by using a 100 W mercury arc lamp (HBO 100) through a bandpass filter (Ex BP/565/30) (Carl Zeiss) before obtaining images. For ATP imaging, signals were monitored by an LSM 5 Pascal confocal laser-scanning microscope equipped with an argon laser (Ex 488, Ex 458) (Carl Zeiss) with a resolution of 1024 × 1024 pixels using a 40× objective lens. Axonal segments containing terminal ends were subjected to imaging analysis.

### In vivo electroporation

In vivo electroporation of mouse cerebella was performed as previously described (20, 33). To visualize mitochondria and axons, 2 μg of tdTomato-Mito-7 (Addgene #58115) (36) and 2 μg of pEGFP-C1 (Takara Bio, Shiga, Japan), together with 0.5 μg of expression plasmid for bcl-xl, were utilized in 10 animals. P6 mice were anesthetized by hypothermia, and DNA in 0.3% fast green/PBS was injected into the surface of the cerebellar cortex with a microsyringe (Hamilton, Reno, NV). Animals were exposed to square electric pulses (4 pulses of 130–140 V for 50 ms with 950 ms intervals) by a pulse generator (CUY edit II) using a tweezer-type electrode attached to the head. Four days later (P10), mice were fixed by cardiac perfusion using 4% PFA in PBS under anesthesia. Cerebella were soaked overnight in a solution containing 30% sucrose in PBS at 4 °C and mounted in OTC compound (Sakura-Finetek, Alphen aan den Rijn, The Netherlands). Coronal sections (60 μm thick) were prepared by a cryostat microtome (CM1850, Leica, Wetzlar, Germany). After staining nuclei with Hoechst 33258 (Sigma-Aldrich), z-stack images were obtained using ApoTome 2 (Carl Zeiss) with a resolution of 1388 × 1040 pixels by a 20× objective lens. Unlike in vitro culture, few images of long axonal segments containing terminals were obtained; thus, in vivo analysis was performed without being limited to such regions.

### Thy1-mitoYFP mice

To generate *Thy1-mitoYFP* mice, we obtained a DNA vector used to generate *Thy1-mitoCFP* mice (Jackson Laboratory, Bar Harbor, ME), which express cyan fluorescent protein (CFP) fused with human cytochrome c oxidase under the regulatory element of the mouse *Thy1* gene. After replacing the *CFP* gene with the *YFP* gene, the constructed vector was injected into fertilized donor mouse eggs from C57/B6J mice. The generated mice expressed YFP protein targeting mitochondria exclusively in neuronal cells, including RGCs (Fig. S2).

To obtain mitochondrial signals from retinal whole mounts of *Thy1-mitoYFP* mice, five 12-week-old male mice were terminally anaesthetized and perfusion-fixed with 4% PFA in PBS following a PBS flush. We enucleated the eyeballs, and the cornea was cut around the edge to remove the lenses and vitreous bodies. We then gently peeled out the retinas from the choroid and the sclera. After washing with PBS, the retinas were placed in 4% PFA in PBS and fixed for another hour. The retinas were rinsed with PBS and flattened by making four radial cuts. Afterward, they were mounted on slide glass and coverslipped using (Lab Vision PermaFluo (Thermo Fisher Scientific Inc.). We observed YFP signals in the whole-mount retinas with a FV1200 confocal microscope (Olympus, Tokyo, Japan) and acquired the images with a resolution of 1024 × 1024 pixels using a 40× objective lens at a 0.5 μm step at the peripheral area approximately 100 μm from the ora serrata. We stacked the images to show the very surface of the retina and retinal ganglion cell layer and processed the stacked images to analyze the position of mitochondria within the axons.

### Data analysis

Images of axons were analyzed using AxioVison software (Carl Zeiss) and subjected to analyses using ImageJ software (National Institute of Health, Bethesda, MD). To analyze the distribution of mitochondria, the place for imaging was decided based on tubulin staining without observing mitochondrial signals. Quantification was conducted automatically by particle analysis using ImageJ. Mitochondria were deemed to be stationary when the maximum change in position during observation was less than 5 μm on kymographs or intermittent images. In the latter case, even if the position did not change, those whose area difference was more than twice were determined to be different mitochondria.

The *Iδ*-index was calculated from the following equation (1), where *q* is the number of parcels and *x*_*i*_ is the mitochondrial number in the parcel.

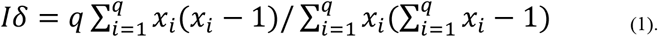

The fitting curve of stationary mitochondria in Fig. 1F was obtained from the one-phase decay equation, where *M*_*s*_ is the ratio of stationary mitochondria that undergo turnover, *M*_*0*_ is the immobile mitochondrial ratio, and *λ* is the decay rate. The half-life (*t*_1/2_) was calculated as follows (3):

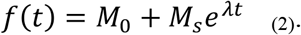

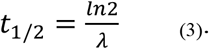

In bar and line graphs, data are expressed as the mean ± 95% confidence interval (CI). In the text, values are presented as the mean ± S.E.M. We used Welch’s t-test for statistical analysis unless otherwise stated. For multiple comparisons, we used Tukey’s test. The levels of significance are denoted as follows: *p < 0.05, **p < 0.01, ***p < 0.001.

## ACKNOWLEDGEMENTS

We are grateful to Dr. Hiroki Takada for the data analysis advice. The GW1-PercevalHR was a gift from Gary Yellen. The mCherry-Mito-7, tdTomato-Mito-7 and mWasabi-Mito-7 were gifts from Michael Davidson. This work was supported by a JSPS KAKENHI Grant Number 20K06889 (Y.K.) and grants from the Life Science Innovation Center of the University of Fukui (Y.K., S.M. and T.T.).

## AUTHIR CONTRIBUTIONS

Author contributions: N. M and Y.K. designed the research; N.M., I.H., T.T., and Y.K. performed the research; T.T., S.M., and M.I. contributed new reagents/analytic tools; N.M., I.H., T.M. and Y.K. analyzed the data; and Y.K. wrote the paper with the contribution of T.T.

## REFERENCES

1. J. S. Kang et al., Docking of axonal mitochondria by syntaphilin controls their mobility and affects short-term facilitation. Cell 132, 137–148 (2008).

2. J. Courchet et al., Terminal axon branching is regulated by the LKB1-NUAK1 kinase pathway via presynaptic mitochondrial capture. Cell 153, 1510–1525 (2013).

3. M. Spillane, A. Ketschek, T. T. Merianda, J. L. Twiss, G. Gallo, Mitochondria coordinate sites of axon branching through localized intra-axonal protein synthesis. Cell Rep 5, 1564–1575 (2013).

4. T. L. Lewis, S. K. Kwon, A. Lee, R. Shaw, F. Polleux, MFF-dependent mitochondrial fission regulates presynaptic release and axon branching by limiting axonal mitochondria size. Nat Commun 9, 5008 (2018).

5. G. M. Smith, G. Gallo, The role of mitochondria in axon development and regeneration. Dev Neurobiol 78, 221–237 (2018).

6. M. J. Devine, J. T. Kittler, Mitochondria at the neuronal presynapse in health and disease. Nat Rev Neurosci 19, 63–80 (2018).

7. R. L. Morris, P. J. Hollenbeck, The regulation of bidirectional mitochondrial transport is coordinated with axonal outgrowth. J Cell Sci 104 (Pt 3), 917–927 (1993).

8. N. Ohno et al., Myelination and axonal electrical activity modulate the distribution and motility of mitochondria at CNS nodes of Ranvier. J Neurosci 31, 7249–7258 (2011).

9. T. L. Lewis, G. F. Turi, S. K. Kwon, A. Losonczy, F. Polleux, Progressive Decrease of Mitochondrial Motility during Maturation of Cortical Axons In Vitro and In Vivo. Curr Biol 26, 2602–2608 (2016).

10. P. J. Hollenbeck, W. M. Saxton, The axonal transport of mitochondria. J Cell Sci 118, 5411–5419 (2005).

11. N. Hirokawa, S. Niwa, Y. Tanaka, Molecular motors in neurons: transport mechanisms and roles in brain function, development, and disease. Neuron 68, 610–638 (2010).

12. K. Obashi, S. Okabe, Regulation of mitochondrial dynamics and distribution by synapse position and neuronal activity in the axon. Eur J Neurosci 38, 2350–2363 (2013).

13. Z. H. Sheng, Q. Cai, Mitochondrial transport in neurons: impact on synaptic homeostasis and neurodegeneration. Nat Rev Neurosci 13, 77–93 (2012).

14. Y. Takihara et al., In vivo imaging of axonal transport of mitochondria in the diseased and aged mammalian CNS. Proc Natl Acad Sci U S A 112, 10515–10520 (2015).

15. N. Ohno et al., Mitochondrial immobilization mediated by syntaphilin facilitates survival of demyelinated axons. Proc Natl Acad Sci U S A 111, 9953–9958 (2014).

16. K. E. Miller, M. P. Sheetz, Axonal mitochondrial transport and potential are correlated. J Cell Sci 117, 2791–2804 (2004).

17. T. Yamada et al., Sumoylated MEF2A coordinately eliminates orphan presynaptic sites and promotes maturation of presynaptic boutons. J Neurosci 33, 4726–4740 (2013).

18. A. Gutnick, M. R. Banghart, E. R. West, T. L. Schwarz, The light-sensitive dimerizer zapalog reveals distinct modes of immobilization for axonal mitochondria. Nat Cell Biol 21, 768–777 (2019).

19. M. Morishita, Measuring of Dispersion of Individuals and Analysis of the Distribution Patterns. Mem. Fac. Sci., Kyushu Univ., Ser. E (Biol.) 2, 215–235 (1959).

20. Y. Konishi, J. Stegmuller, T. Matsuda, S. Bonni, A. Bonni, Cdh1-APC controls axonal growth and patterning in the mammalian brain. Science 303, 1026–1030 (2004).

21. A. Ito-Ishida et al., Presynaptically released Cbln1 induces dynamic axonal structural changes by interacting with GluD2 during cerebellar synapse formation. Neuron 76, 549–564 (2012).

22. M. Tantama, J. R. Martínez-François, R. Mongeon, G. Yellen, Imaging energy status in live cells with a fluorescent biosensor of the intracellular ATP-to-ADP ratio. Nat Commun 4, 2550 (2013).

23. A. Oruganty-Das, T. Ng, T. Udagawa, E. L. Goh, J. D. Richter, Translational control of mitochondrial energy production mediates neuron morphogenesis. Cell Metab 16, 789–800 (2012).

24. K. Fukumitsu et al., Synergistic action of dendritic mitochondria and creatine kinase maintains ATP homeostasis and actin dynamics in growing neuronal dendrites. J Neurosci 35, 5707–5723 (2015).

25. M. E. Bulina et al., A genetically encoded photosensitizer. Nat Biotechnol 24, 95–99 (2006).

26. A. Vaarmann et al., Mitochondrial biogenesis is required for axonal growth. Development 143, 1981–1992 (2016).

27. K. T. Chang, R. F. Niescier, K. T. Min, Mitochondrial matrix Ca2+ as an intrinsic signal regulating mitochondrial motility in axons. Proc Natl Acad Sci U S A 108, 15456–15461 (2011).

28. Y. Chen, Z. H. Sheng, Kinesin-1-syntaphilin coupling mediates activity-dependent regulation of axonal mitochondrial transport. J Cell Biol 202, 351–364 (2013).

29. S. Lee, W. Wang, J. Hwang, U. Namgung, K. T. Min, Increased ER-mitochondria tethering promotes axon regeneration. Proc Natl Acad Sci U S A 116, 16074–16079 (2019).

30. B. Zhou et al., Facilitation of axon regeneration by enhancing mitochondrial transport and rescuing energy deficits. J Cell Biol 214, 103–119 (2016).

31. D. Zala et al., Vesicular glycolysis provides on-board energy for fast axonal transport. Cell 152, 479–491 (2013).

32. T. Seno et al., Kinesin-1 sorting in axons controls the differential retraction of arbor terminals. J Cell Sci 129, 3499–3510 (2016).

33. Y. Inami, M. Omura, K. Kubota, Y. Konishi, Inhibition of glycogen synthase kinase-3 reduces extension of the axonal leading process by destabilizing microtubules in cerebellar granule neurons. Brain Res 1690, 51–60 (2018).

34. S. G. Olenych, N. S. Claxton, G. K. Ottenberg, M. W. Davidson, The fluorescent protein color palette. Curr Protoc Cell Biol Chapter 21, Unit 21.25 (2007).

35. M. A. Rizzo, M. W. Davidson, D. W. Piston, Fluorescent protein tracking and detection: fluorescent protein structure and color variants. Cold Spring Harb Protoc 2009, pdb.top63 (2009).

36. H. W. Ai, K. L. Hazelwood, M. W. Davidson, R. E. Campbell, Fluorescent protein FRET pairs for ratiometric imaging of dual biosensors. Nat Methods 5, 401–403 (2008).

