## supplemental figures, methods for "Intermitochondrial signaling regulates the uniform distribution of stationary mitochondria in axons"

Supporting Information

### SI Materials and Methods

*Culture of RGC neurons*

Primary culture of rat RGCs were prepared from P3 Sprague-Dawley rats (Charles River Japan, Yokohama, Japan) by a two-step immunopanning method using anti-macrophage antibodies and anti-thy1.1 antibodies as previously described (1, 2). Neurons were plated on glass bottom plates pretreated with Poly-D-lysine at 2500 cells/cm$^2$, and maintained in Neurobasal Medium without phenol red, containing B27 supplement (Thermo Fisher Scientific Inc.), brain-derived neurotrophic factor (50 ng/mL), ciliary neurotrophic factor (50 ng/mL), basic fibroblast growth factor (50 ng/mL), forskolin (10 μM), glutamine (1 mM), insulin (5 μg/mL), sodium pyruvate (1 mM), sodium selenite (40 ng/mL), transferrin (100 μg/mL), triiodothyronine (30 ng/mL), triiodothyronine, penicillin (1 mg/ml) and streptomycin (1 mg/ml).

*Imaging and analysis of mitochondria in RGC axons in vitro*

To label mitochondria, 3.2 μg of tdTomato-Mito plasmid, 2 μl of Lipofectamine LTX reagent and 2 μl of PLUS reagent (Thermo Fisher scientific Inc.) were mixed in 115μl of Opti-MEM (Thermo Fisher scientific Inc), and 16 μl of mixture was added for one 35 mm glass bottom dish at 5 DIV. Axonal mitochondria in RGCs were monitored by using an FV10i (Olympus) at 8 DIV. To determine the positions of stationary mitochondria, time-lapse images were converted to the kymograph by using ImageJ. Compartmental analysis was performed in the same procedure as for CGN.

### SI References


1. B. A. Barres, B. E. Silverstein, D. P. Corey, L. L. Chun, Immunological, morphological, and electrophysiological variation among retinal ganglion cells purified by panning. *Neuron* 1, 791-803 (1988).
2. S. Miyake, Y. Takihara, S. Yokota, Y. Takamura, M. Inatani, Effect of Microtubule

Disruption on Dynamics of Acidic Organelles in the Axons of Primary Cultured Retinal Ganglion Cells. *Curr Eye Res* 43, 77-83 (2018).

### SI Figure Legends

Fig. S1. Distribution of stationary along RGC axons in vitro.

(A) Representative kymograph obtained from an 8 DIV RGC axons expressing tdTomato-Mito. Scale bar indicates 20 μm. (B) Iδ-index profiles of axonal mitochondrial spots obtained from cultured RGC axons. Values represent mean ± 95%CI. (C) Frequency of observed mitochondrial spots (solid line) versus Poisson distribution (dotted line) at 20 μm compartment size indicated that mitochondrial spots distribution is significantly different from random distribution (*p < 0.05, $\chi^2$ analysis). Data was obtained from n = 12 axons.

Fig. S2. Visualization of mitochondria in RGC in retina.

Flat mount of retina prepared from *Thy1-mitoYFP* transgenic mouse at 12 weeks old. Mitochondria in RGC neurons were visualized. The arrow indicates the direction of optic disc (OD). Scale bar indicates 500 μm.

Fig. S3. Effect of ATP depletion on the signal of PercevelHR. (A) CGNs that express PercevalHR were treated with 1 μM of olygomycin (OM) or 50 mM of 2-deoxyglucose (2DG). Signals for ATP (Ex 488; green) and isosbestic point (Ex 458; red) at 1 to 2 hrs after treatment were shown. Scale bar indicates 50 μm. (B) Olygomycin treatment significantly reduced the Ex 488/Ex 458 ratio. 2-deoxyglucose slightly reduced the signal. The value for each experiment was calculated from the 5 neurons with the strongest signal. **p < 0.01, ***p < 0.001, n = 4 experiments, Tukey's test. Values represent mean ± 95%CI.

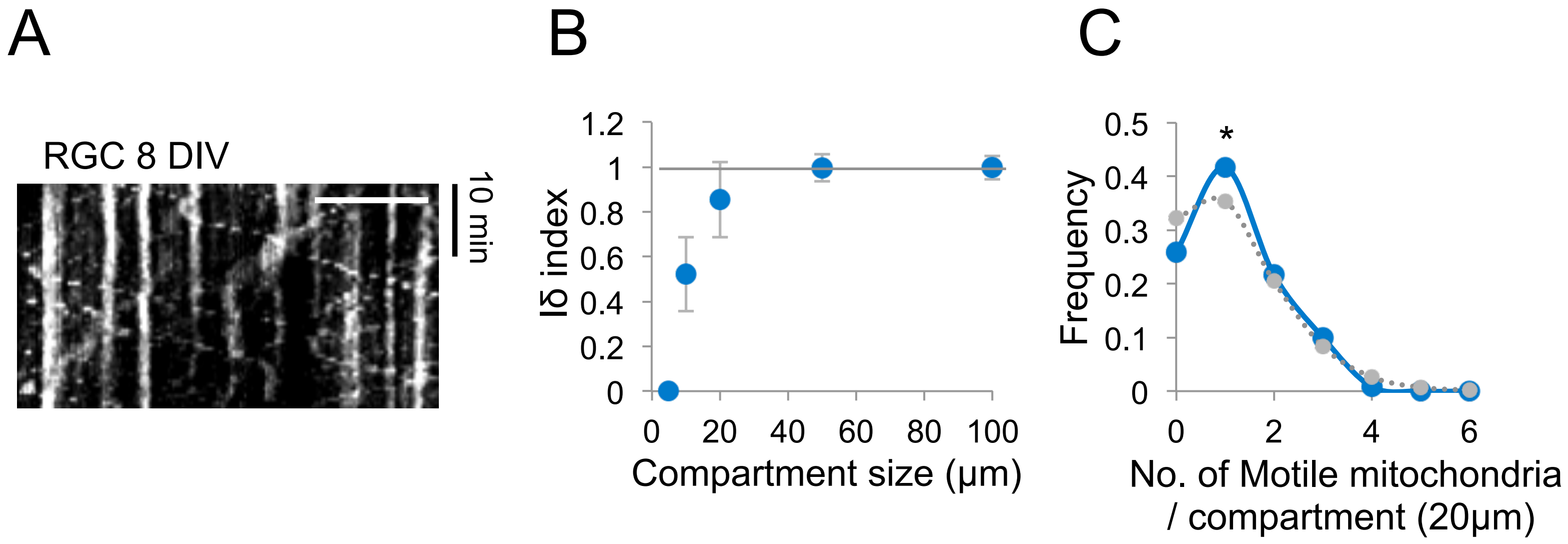
A
RGC 8 DIV
10 min
B
1.2
1
0.8
0.6
0.4
0.2
0
Iδ index
0
20
40
60
80
100
Compartment size (μm)
C
*
0.5
0.4
0.3
0.2
0.1
0
Frequency
0
2
4
6
No. of Motile mitochondria
/ compartment (20μm)

Fig. S1

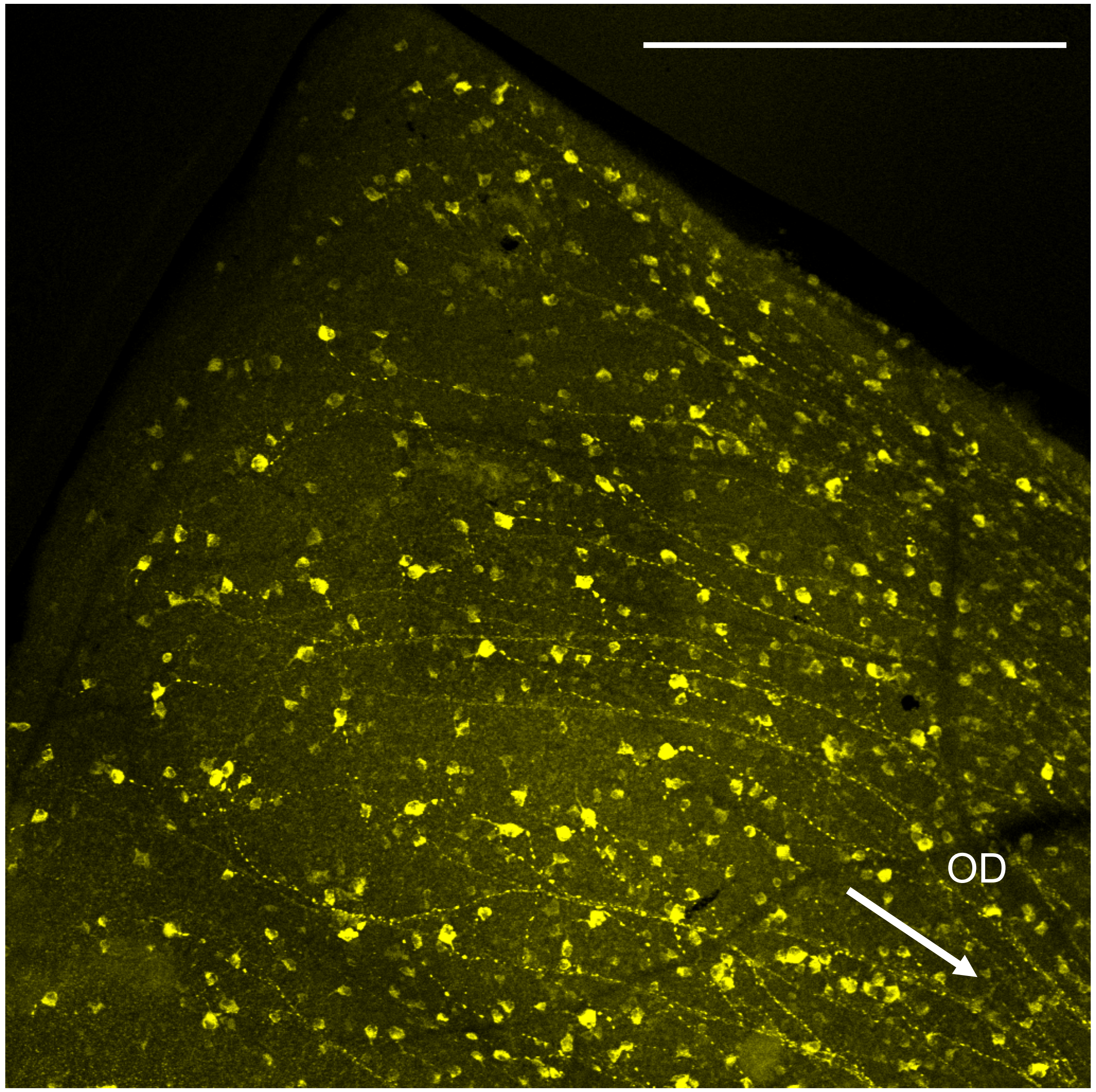


Fig. S2

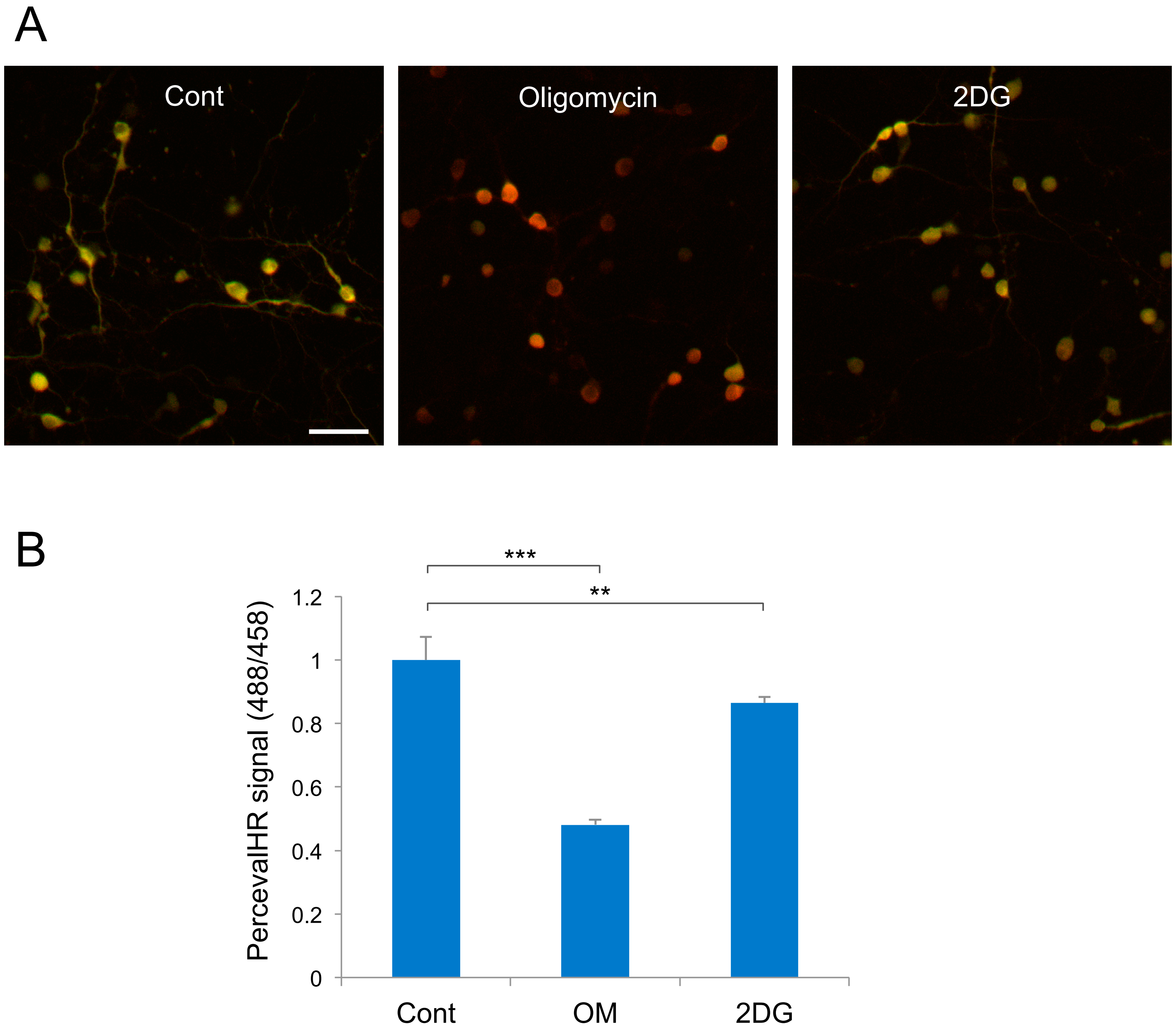
A
Cont
Oligomycin
2DG
B
***
**
PercevalHR signal (488/458)
1.2
1
0.8
0.6
0.4
0.2
0
Cont
OM
2DG

Fig. S3
